# *In situ* molecular architecture of the mammalian sperm nuclear vacuole

**DOI:** 10.64898/2026.05.18.725999

**Authors:** Julian R. Braxton, Songrong Qu, Will M. Skinner, Momoko Shiozaki, Nikki Jean, Rui Yan, Xiaowei Zhao, Zhiheng Yu, Polina V. Lishko, Zhen Chen

## Abstract

Direct identification of macromolecular complexes in their native context remains a major barrier to unbiased biological discovery. This challenge is particularly acute in mammalian sperm nuclei, in which condensed chromatin is interspersed with poorly understood phase-separated compartments termed nuclear vacuoles. Vacuoles are associated with reduced fertilization efficiency, yet their composition remains unclear. Here we combine high-resolution *in situ* cryo-electron tomography (cryo-ET) with AlphaFold docking to identify vacuole components as proteasomes, the proteasome activator PA200, and ferritin. *In situ* structures at resolutions up to 3.8 Å reveal distinct proteasome-PA200 associations and gating states, consistent with a stepwise activation mechanism. Ferritin assemblies exhibit heterogeneous mineralization states and directly contact chromatin. Together, these findings establish the molecular organization of sperm nuclear vacuoles and implicate protein turnover and metal homeostasis in shaping the nuclear landscape, while demonstrating the power of *in situ* cryo-ET to resolve protein identity and conformational dynamics in native cellular environments.

## Introduction

Electron microscopy has long provided ultrastructural views of cellular organization^1,2^, but chemical fixation and staining in conventional workflows obscure molecular detail, limiting the identification and analysis of macromolecules in their native states. Cryo-electron tomography (cryo-ET) has emerged as a high-resolution imaging technique for unfixed, unstained biological material in a frozen-hydrated state^3,4^. Critically, this technology has the potential to enable high-resolution structure determination *in situ*, enabling de novo identification of molecular components and the generation of hypotheses about their cellular functions^5^. This capability is particularly needed in the study of mammalian sperm nuclei, which represent an extreme chromatin state in which the genome is compacted to less than 5% of the volume of a somatic nucleus^6^ through histone replacement by sperm-specific DNA-binding proteins called protamines^7–9^. Despite this dramatic condensation, sperm nuclei contain enigmatic phase-separated compartments known as nuclear vacuoles^10^. The presence of vacuoles correlates with reduced motility and fertilization efficiency^11,12^, yet their composition remains unknown. Here we used *in situ* cryo-ET to resolve macromolecular assemblies directly within human sperm nuclear vacuoles at near-atomic resolution, identifying them as proteasome complexes, a subset of which is bound to the ubiquitin-independent activator PA200, and ferritin. These findings uncover a previously unrecognized molecular organization within the sperm nucleus and implicate proteolysis and metal handling in the development and function of this specialized organelle.

## Results

### Cryo-ET of sperm nuclear vacuoles reveals distinct protein complexes

As human sperm nuclei are too thick to be directly imaged by transmission electron microscopy, we generated lamellae of sperm nuclei by cryo-FIB milling^13^ and collected tilt series on electron-lucent regions corresponding to vacuoles (Figure S1A). Inspection of reconstructed tomograms revealed that vacuoles are membrane-less, phase-separated compartments of variable dimensions (∼15 nm to > 1 µm in diameter) interspersed among dense material likely corresponding to condensed chromatin (Figure 1A, S1B). The non-spherical appearance and rough edges of vacuoles suggest non-liquid behaviors of condensed chromatin. We also observed distinct nuclear morphologies, in which vacuoles formed interconnected networks among condensed chromatin, rather than discrete structures (Figure S1C). Notably, numerous protein complexes of varying size (∼120-280 Å in length) were identified in vacuoles (Figure 1A, insets). The high contrast of these complexes against the vacuole background indicated the absence of chromatin or other proteins from these compartments. Three-dimensional rendering of identified particles in a representative tomogram (Figure 1A) revealed a densely packed arrangement in the vacuole (Figure 1B) and no well-defined macromolecules were identified in sperm chromatin, likely hindered by the strong signal of condensed chromatin. Next, to determine whether sperm nuclear vacuoles and their resident proteins are unique to human sperm, we performed our cryo-ET workflow on mouse sperm. Inspection of reconstructed tomograms revealed vacuoles of similar appearance with internal protein complexes in mouse sperm, although with decreased size and frequency (Figure S1D). These results suggest that nuclear vacuoles enclosing discrete protein complexes are a conserved feature of mammalian sperm.

**Figure 1.**
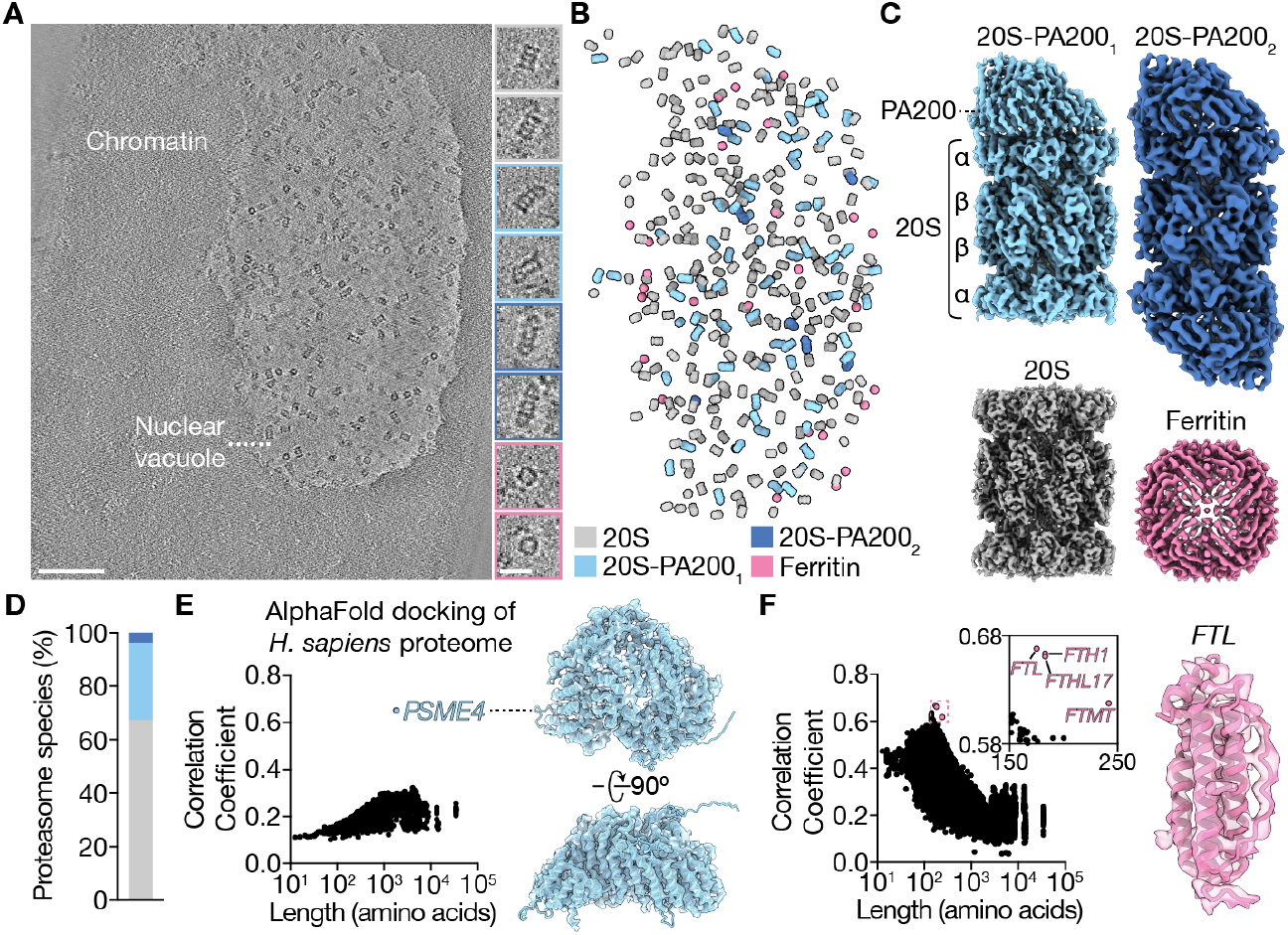
Cryo-ET of human sperm nuclear vacuoles reveals distinct protein complexes. **(A)** Left, z-slice of a representative human sperm nucleus tomogram (denoised). Right, gallery of individual particles identified in human sperm nuclear vacuoles (central z-slices). Colored outlines correspond to assigned identity in **(B, C). (B)** Three-dimensional rendering of identified particles after sub-tomogram averaging in a 40 nm thick slice of the tomogram in **(A)**, colored by identity. **(C)** Sub-tomogram averages of the four identified species in human sperm nuclear vacuoles (filtered maps). **(D)** Relative abundance of all identified 20S, 20S-PA200_1_, and 20S-PA200_2_ particles (*n* = 111 tomograms). **(E)** Left, docking of the human AlphaFold proteome into the additional density on 20S proteasome complexes. Right, orthogonal views of the best-fitting AlphaFold model, PA200 (gene name *PSME4*), overlaid with density used for docking. **(F)** Left, docking of the human AlphaFold proteome into the asymmetric unit of the ferritin map. Ferritin homologs are colored pink and labeled with gene names. Right, view of the best-fitting AlphaFold model, *FTL*, overlaid with the density used for docking. Scale bars, 100 nm (**A**, left), 20 nm (**A**, right inset).

We next employed sub-tomogram averaging to reconstruct the macromolecular complexes in human sperm vacuoles, and AlphaFold^14,15^ docking to determine their identities. Three-dimensional classification and multi-particle refinement in M^16^, in which all species were jointly refined to improve alignment quality, yielded four reconstructions with global resolutions ranging from 3.8-7.8 Å (Figure 1C, S2, S3A-D, Table S1). The quality of these maps, all with resolved secondary structure, enabled the unambiguous identification of three proteasome complexes, characterized by four stacked rings of seven subunits each. One proteasome species corresponded to the 20S core particle, and comprised ∼67% of all identified proteasomes (Figure 1D). The other two proteasome species contained one or two additional densities contacting the α ring, comprising ∼29% and ∼4% of proteasomes, respectively. The fourth species was identified as a hollow, octahedrally symmetric sphere ∼120 Å in diameter. We performed unbiased docking of the human AlphaFold2 proteome into the additional density on proteasomes and the asymmetric unit of the spherical complex (Methods). The map-model correlation coefficient for the best-scoring pose was plotted for 23,577 human protein structures (Figure 1E,F). This analysis revealed an unambiguous identity for the proteasome-associated density (correlation ∼0.65; mean +/-standard deviation, 0.2 +/-0.04): the ubiquitin-independent proteasome activator PA200 (Figure S4). Docking of PA200 into the map revealed matching secondary and tertiary structures throughout the 211 kDa protein, with discrepancies only for an N-terminal extension and several internal loops (Figure 1E, right). Docking results for the asymmetric unit of the spherical map revealed top candidates containing four-helical bundles (19 of the top 20, Figure S5). The proteins with the three highest scores were all homologs of ferritin, a typically cytosolic oligomeric iron storage protein^17^ not previously reported in sperm nuclei (Figure 1F, right). While many models fit the four-helix density well, only ferritin accounted for an ordered loop at the periphery of the structure (Figure S5) and a fifth, smaller α-helix (helix E). Moreover, unlike other top hits, ferritin oligomerizes into highly symmetric nanocages^18^ of similar appearance to those observed *in situ*, while most false positives (15 of the 16 remaining non-ferritin proteins in the top 20) were membrane proteins that would not oligomerize into a nanocage. These results strongly suggest that ferritin is the correct candidate. Taken together, our results indicate that the primary macromolecular components of sperm nuclear vacuoles are proteasomes (20S and 20S-PA200) and ferritin complexes.

### High-resolution *in situ* proteasome structures enable paralog and isoform determination

To analyze the composition and conformation of proteasomes in sperm vacuoles, we reconstructed high-resolution maps of 20S and 20S-PA200 complexes using the *in situ* data. During classification we observed PA200 density of varying strength relative to the 20S, indicating conformational heterogeneity (Figure S6). After selecting particles with strong PA200 density, refinement resulted in a 20S-PA200 consensus map at 4.5 Å and a 20S map at 3.8 Å, in which side chains of large residues were resolved (Figure 2A-C, S2, S3A,B, Table S1). These high-resolution maps enabled building of atomic models, which revealed structural similarity to 20S and 20S-PA200 structures determined *in vitro* (Cα root mean squared deviations of 1.8 Å and 1.7 Å, respectively)^19,20^. In both structures, the 20S was identified as a hollow 28-mer cylinder composed of α and β subunits in an α1-7,β1-7,β1-7,α1-7 configuration. Interior proteolytic sites of β subunits are shielded from the cellular milieu by N-terminal extensions of α subunits, which form a gate that is only opened by the association of proteasome activators. Importantly, strong map-model correspondence and resolved asymmetric features such as the closed α gate formed by α2-4 and the C-terminal extension of β2^21^ demonstrated the correct alignment of the 20S structure despite its pseudo-*D*7 symmetry (Figure 2D). In the 20S-PA200 structure, PA200 adopted an asymmetric arrangement on the 20S α ring, contacting all seven subunits but largely positioned over α1-α5 (Figure 2A, bottom left). The α gate proximal to PA200 (the *cis* gate) was open, exposing the interior catalytic residues of the β subunits. In contrast, the *trans* α gate, and those of the 20S complex, were closed by α subunit N-terminal extensions. These results indicate that binding of PA200 opens the 20S proteasome gate in native sperm vacuoles.

**Figure 2.**
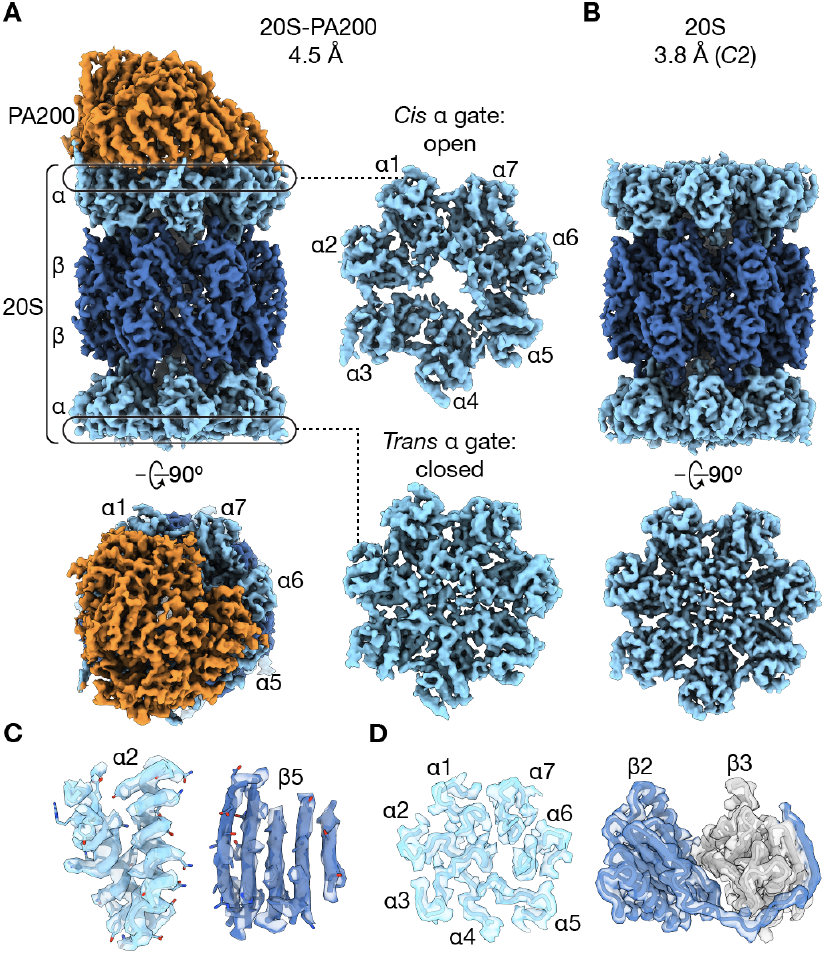
*In situ* analysis enables proteasome structure determination at near-atomic resolution. **(A)** Left, views of the 20S-PA200 consensus map (filtered). 20S α and β subunits are colored light and dark blue, respectively; PA200 is colored orange. Right, views of the *cis* (contacting PA200) and *trans* (not contacting PA200) α gates. **(B)** Views of the filtered 20S map, colored as in **(A). (C)** High-resolution features of the 20S map. Segmented maps (locally filtered, transparent surfaces) and corresponding models are shown. **(D)** Asymmetric features of the 20S map. Segmented maps (filtered, transparent surfaces) and corresponding models are shown. Left, α gate. Right, β2 subunit with C-terminal extension (blue) and β3 subunit (gray).

We next analyzed the subunit composition of the *in situ* 20S proteasome. Prior mass spectrometry and Western blotting data^22,23^ suggested that α4, β2, and β5 are replaced in proteasomes found in sperm by a testis-specific α4 paralog (α4s), and the immunoproteasome subunits β2i and β5i. However, proteasome composition(s) in nuclear vacuoles are unknown. Our high-resolution 20S density map enabled the identification of side chains belonging to the constitutive β2 and β5 in sperm nuclear proteasomes (Figure S7A,B). In contrast, the majority of the density at α4 could be modeled similarly by α4 and α4s upon visual inspection (Figure S7C). Surprisingly, although a six-residue extension in full-length α4s was inconsistent with the density, a reported alternatively-spliced isoform lacking these residues also fit the density well (Figure S7D)^24^. Rosetta modeling of the alternatively-spliced α4s isoform indicated a better fit than for the constitutive α4, suggesting that this isoform is the main component in sperm proteosomes in nuclear vacuoles (Figure S7E and Methods). These analyses demonstrate that our cryo-ET workflow can achieve high-resolution reconstructions and directly read side chain information to distinguish close paralogs and isoforms inside cells.

### Direct visualization of ferritin mineralization and chromatin contacts

We next analyzed ferritin structures and localization in vacuoles. We employed three-dimensional classification to maximize the quality of the particle set, resulting in 1,635 particles from 92 tomograms that were refined to 6.1 Å with octahedral symmetry (Figure 3A, S2, S3D, Table S1). This reconstruction agreed with previously determined structures of intact ferritin oligomers^25^, in which 24 protomers enclose a central cavity inside which reactive or toxic metals can be sequestered from the cellular environment^26^ (termed holo-ferritin, in contrast to metal-free apoferritin). Additionally, we observed density for the C-termini of ferritin protomers, which extend from helix E and point toward the center of the nanocage from all *C*4 vertices^27^ (Figure 3A, “*”). We did not observe any additional metal density in our consensus reconstruction; however, such particles may have been excluded by auto-picking and classification algorithms due to the stronger signals of heavy elements compared to light element-based proteins. Manual inspection of tomograms, however, revealed nanocages with strongly scattering interior densities of varying appearance that likely correspond to metal-rich deposits (Figure 3B). These deposits ranged from ∼1-7 nm in diameter and in some views appeared localized to the periphery of the cage interior. Reconstructions of apo- and holoferritin using manually-picked particles resembled the consensus structure, but the holoferritin map featured strong central density (Figure 3C, S2). The voxel intensity of this density is approximately six-fold greater than that of the protein shell, consistent with a composition enriched in strongly scattering elements such as iron^28^ (Figure 3D). Connections of the central density to the *C*4 ferritin vertices suggested that the metal-ferritin interaction was mediated by helix E and/or its C-terminal extension (Figure 3C, right inset). These results demonstrate that high-resolution *in situ* cryo-ET can reveal molecular identities of even minor species (∼15 particles per tomogram, Figure S8) and suggest ferritin plays a role in metal homeostasis in nuclear vacuoles.

**Figure 3.**
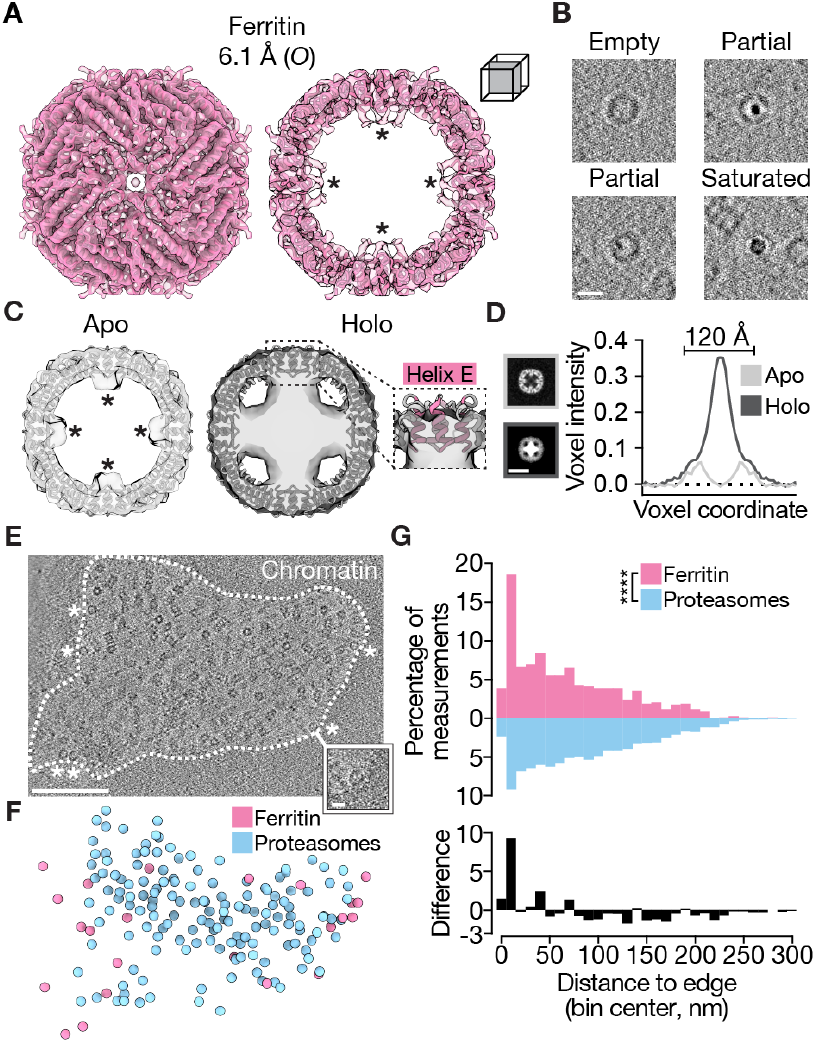
Direct visualization of ferritin mineralization and chromatin contacts. **(A)** Ferritin cryo-ET reconstruction (transparent, locally filtered) and fitted atomic model of human ferritin heavy chain (PDB 7A6A). Right, central slice of structure. Density for the C-terminal extensions of helix E is indicated (*). **(B)** Gallery of ferritin particles from human sperm nuclear vacuole tomograms (central z-slices), showing variable interior density. **(C)** Central slices of apo- and holoferritin reconstructions, overlaid with an atomic model of human ferritin heavy chain (PDB 7A6A). Density for the C-terminal extensions of helix E is indicated in the apoferritin map (*). Putative ferritin-metal contact in holoferritin is shown as an enlarged view. **(D)** Left, central slices, and right, edimensional voxel intensities of apoand holoferritin maps in **(C). (E)** Tomogram z-slice showing putative ferritin-chromatin contacts. Ferritin particles adjacent to chromatin are indicated (*), and an enlarged view of such a particle is outlined in white. The vacuole boundary is shown as a dashed line. **(F)** Three-dimensional rendering of ferritin (pink spheres) and proteasomes (blue spheres) in the nuclear vacuole shown in **(E). (G)** Histogram of ferritin and proteasome distances to vacuole edge (median distances, 55 and 78 nm, respectively), and calculated difference between these values (*n* = 10 tomograms). ****, p < 0.0001, Welch’s two-tailed t-test. Scale bars, 10 nm (**B, D, E**, inset), 100 nm (**E**, tomogram).

When inspecting ferritin particles in tomograms, we noticed that many were localized to the vacuole periphery, and appeared to directly contact chromatin (Figure 3E,F, 1A,B). To determine whether ferritin particles were non-uniformly distributed in vacuoles, we measured the distances of ferritin and proteasomes to the vacuole edge in tomograms in which the xy boundary of the vacuole was fully visible (Methods). Notably, the edge distance distributions for ferritin and proteasomes were significantly different (p < 0.0001, Methods), and a large population of ferritin was localized within 6 nm (ferritin diameter: ∼12 nm) to the vacuole edge (12% vs 6% for ferritin and proteasomes, respectively (Figure 3G). These results are consistent with a direct ferritin-chromatin interaction *in situ*, as previously reported *in vitro*^29^.

### *In situ* dynamics of 20S-PA200 complexes reveal a stepwise activation mechanism

The heterogeneous appearance of PA200 density observed during the generation of the 20S-PA200 consensus map (Figure S6) prompted us to closely investigate these particles. We classified particles into three populations of similar size but with variable PA200 density and 20S contacts (weak, partial, and full classes, Figure 4A,B); subsequent refinement led to maps with resolutions between 6.2 Å and 7.3 Å (Figure S2, S3E-G, Table S1). The full-bound structure resembled the high-resolution 20S-PA200 consensus structure (Figure 2A). Surprisingly, the gates of the other two PA200-bound structures were closed, a conformation not previously reported *in vitro*^19,20,30^. In 20S-PA200^weak^, the full PA200 density was only observed after Gaussian filtering, indicating substantial conformational variability with respect to the 20S core. Moreover, PA200 contacts in this structure were limited to α5-7 (Figure 4A, bottom), suggesting this may be an initial association interface. PA200 density in 20S-PA200^partial^ and 20S-PA200^full^ was readily apparent, though was weaker at the 20S interface of the partial map (Figure 4A). Indeed, a difference map of the two structures revealed stronger PA200 density proximal to the α ring in the full-bound map, particularly near α3-α5 (Figure 4C), suggesting this may be a later association interface. Decreased PA200 local resolution in the partial-bound map was further suggestive of flexibility and incomplete binding in this state (Figure S3F). Interestingly, the C-terminal α helices of α3 and α4 were also better defined in the full-bound state (Figure 4C), suggesting that they are stabilized upon PA200 binding. These results suggest that complete PA200 binding is required for gate opening.

**Figure 4.**
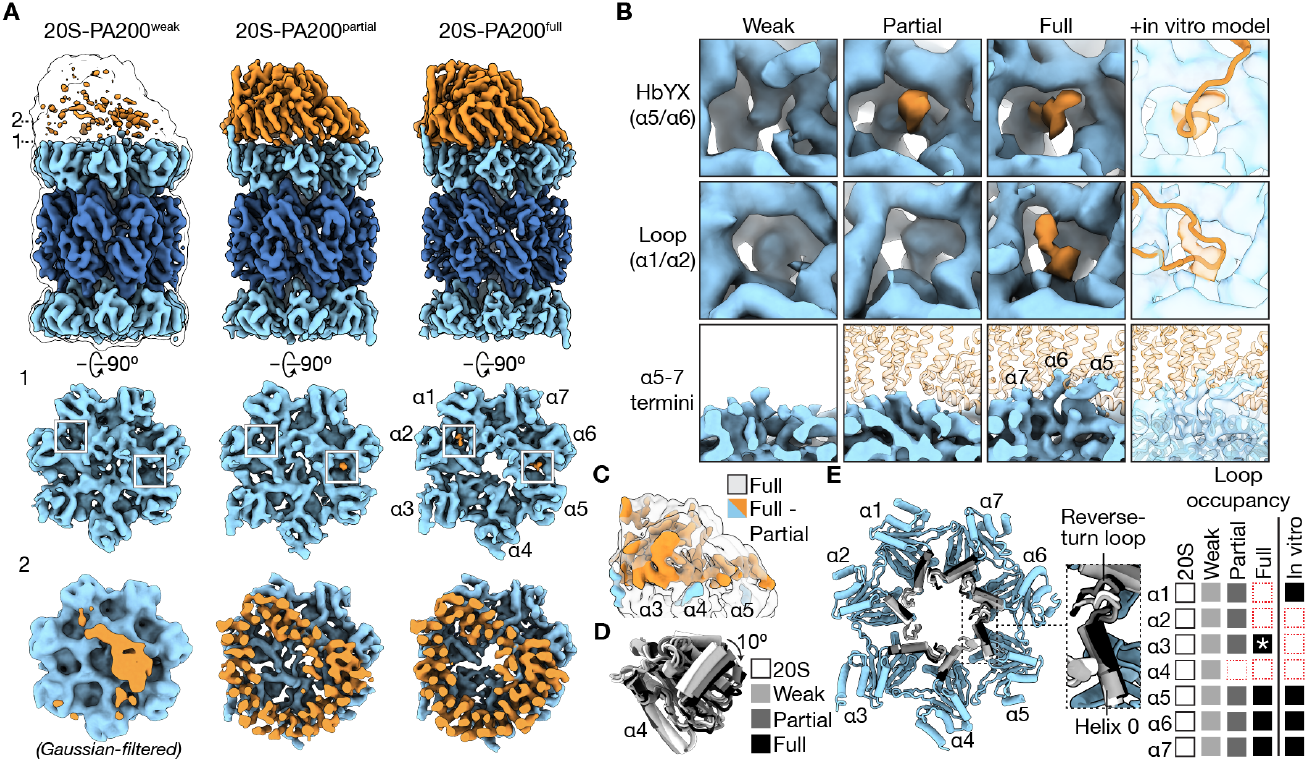
*In situ* analysis of PA200-bound proteasomes reveals a stepwise activation mechanism. **(A)** Top, 20S-PA200 complexes (filtered) from PA200/α ring subclassification, colored as in Figure 2A. 20S-PA200^weak^ is overlaid with a transparent Gaussian-filtered version of the same map, standard deviation 3. Middle, views of the *cis* α gates from the corresponding maps, showing variable PA200 contacts and activation states. α pockets with PA200 contacts in the full state are outlined in all three states. Bottom, views of PA200 density (same clipping plane used for all structures), showing contact in the weak state with only α5-7. The weak map is Gaussian-filtered (standard deviation 3); the partial and full are filtered, as in the above rows. **(B)** Discrete 20S-PA200 contact sites in weak, partial, and full maps (filtered), colored as in **(A)**. Transparent 20S-PA200^full^ views overlaid with relevant regions of an *in vitro* 20S-PA200 structure (PDB 6KWY) are shown at right. **(C)** Difference map of 20S-PA200^full^ and 20S-PA200^partial^ (Gaussian filtered, standard deviation 3, colored as in **(A)**), overlaid with 20S-PA200^full^ (Gaussian filtered, standard deviation 3, transparent). **(D)** Overlay of α4 subunits from α rings of 20S and *cis* 20S-PA200 complexes, aligned by β4, showing outward axial rotation. **(E)** Left, model of *cis* α ring of 20S-PA200^full^ overlaid with helix 0 and reverse-turn loops (where present) from the remaining *in situ* proteasome structures, colored as in **(E)**, showing radial expansion of α rings with increasing PA200 density. An enlarged view of the α5 loops and helix 0 is shown at right. N-terminal gating residues are omitted for clarity. Right, summary of reverse-turn loop occupancy in 20S and 20S-PA200 complexes. Colored squares indicate resolved loops, red dashed outlines indicate disordered loops. A weak but present α3 loop in 20S-PA200^full^ is indicated (*). Loops for structures determined *in vitro* (e.g., PDB 6KWY) are included as a comparison.

We next sought to analyze 20S-PA200 contacts in more detail. Three distinct interactions were made with the α ring in 20S-PA200^full^: insertion of the PA200 C-terminal HbYX (hydrophobic, Tyr, any amino acid) peptide (1841-YYA-1843) into the α5/α6 pocket, insertion of an internal PA200 loop into the α1/α2 pocket (hereafter referred to as the α1/α2 loop), and stabilization of the N-termini of α5-α7 upward into the PA200 cavity (Figure 4B). We observed varying contact patterns in the two closed states: the weak-bound structure had none, the partial-bound structure only had resolved HbYX density. These structural findings indicate that multiple identified interactions are necessary to synergistically open the 20S gate. Notably, *in vitro* structural studies have only revealed PA200 interactions with gate-open proteasomes, demonstrating that *in situ* analysis of a heterogeneous proteasome population can reveal key conformational intermediates.

To quantitatively analyze 20S-PA200 conformational variability, we built models into all three maps (Figure S3E-G, Table S1) and analyzed them alongside the *in situ* 20S model. We observed progressively greater outward axial rotation of the α subunits of the *cis* ring from the weak to partial to full states, to a maximum of ∼10° relative to the 20S alone (Figure 4D, S9A,B). Furthermore, we observed outward movement of helix 0 (H0) and the adjacent reverse-turn loop, structural elements surrounding the gate (Figure 4E). However, distinct patterns of H0 and reverse-turn loop occupancy were observed relative to 20S-PA200 structures determined *in vitro*, a finding that may be explained by *in situ* analysis, the replacement of α4 with α4s, or the retention of heterogenous particles during processing. These findings indicate that 20S-PA200 complexes adopt distinct conformations *in situ*. Interestingly, the largest change in the 20S between the partial-and full-bound states was the outward radial rotation of α4 (Figure S9C), suggesting that this movement may induce gate opening through an extended network of interactions propagated around the α ring. Together, these results suggest that stepwise association of PA200 leads to the gradual displacement of pore-adjacent structural elements, ultimately opening the α gate (Figure 5).

**Figure 5.**
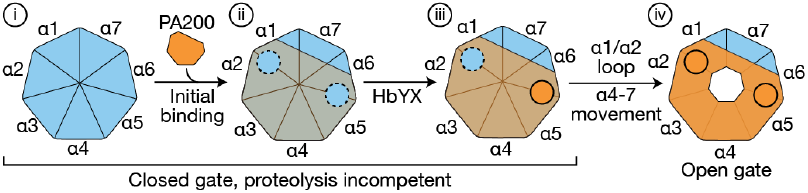
Model of PA200-mediated proteasome activation *in situ*. Stepwise model of PA200-mediated proteasome activation derived from *in situ* cryo-ET structures. i, the 20S α ring (blue) is closed without the association of an activator. ii, initial PA200 binding results in a loose association with the 20S α ring (indicated by an orange transparent overlay) without anchoring into α pockets (black dashed lines). iii, the PA200 C-terminal HbYX peptide docks into the α5/α6 pocket (filled orange circle) and PA200 adopts a more stable conformation on the α ring (more opaque orange), but these interactions are insufficient to induce gate opening. iv, Complete PA200 association, associated with docking of the α1/α2 loop, sequestration of the α5-7 N-terminal extensions, and α4 rotation, results in gate opening, rendering the proteasome competent to degrade substrates.

## Discussion

Methods to directly identify macromolecules in their native context and at high resolution promise to reveal novel aspects of cell biology, yet remain technically challenging. The high-resolution cryo-ET structures obtained here were enabled by recent methodological advances in processing two-dimensional images directly acquired by the microscope, rather than using three-dimensional approximations of these data (subtomograms)^31^. We used a narrow tilt range (±40°) and sparse angular sampling (4°) to increase the signal-to-noise ratio in each tilt image, combined with multi-particle refinement of tilt-series parameters using M^16^. Because the multi-particle approach jointly refines parameters using all particles in a tilt series, rare complexes such as vacuolar ferritin benefit from the accurate alignments contributed by high-resolution reconstructions of proteasomes. Together, these advances enable high-resolution *in situ* imaging of relatively small macromolecules and less abundant particles, thereby broadening the utility of this workflow.

Although proteasomes are highly enriched in the nucleus of various cell types^32^, the functional basis of this enrichment is unclear. Our discovery of abundant PA200-bound proteasomes in sperm nuclear vacuoles suggests that protein turnover is important during nuclear maturation. These proteasomes have been proposed to degrade acetylated histones during spermiogenesis, enabling histone-to-protamine exchange^22^, and to function during meiosis, where the canonical α4 subunit is replaced by the testis-specific α4s^23,33–36^. The presence of α4s-containing proteasomes in mature sperm demonstrates that this specialized form persists into late spermatogenesis. However, the absence of detectable substrate density suggests that vacuolar proteasomes may be a residual population after completing their degradative role. Observations of uncondensed chromatin and interconnected vacuolar regions (Figure S1C) support a model in which vacuoles arise as strong protamine–DNA interactions^37^ compact chromatin into a dense phase while excluding other nuclear components in the corresponding dilute phase. Proteasome and PA200 involvement in chromatin remodeling may extend to other contexts, including the reported role of PA200 in DNA repair^22,38^.

We observed multiple metal morphologies in vacuolar holoferritin complexes. Peripheral small particles (∼1–2 nm) are consistent with current models of mineral nucleation involving acidic residues of ferritin light chain at the interior ferritin surface^39^, and may thus represent early stages of mineralization. Larger aggregates occupying much of the ferritin interior may represent later stages in this process, and density connecting helix E and the ferritin C-terminus to the metal core suggests that this region mediates contact with mineral particles. Similar interactions have been observed in recombinant ferritin light chain^40^, and frameshift mutations that replace helix E and the C-terminus with unrelated sequences impair iron accumulation and cause hereditary diseases termed ferritinopathies^41,42^, supporting the biological relevance of the interaction observed *in situ*. Moreover, ferritin within vacuoles may play a role in protecting germline DNA, as nuclear ferritin^29,43,44^ has been shown to protect genomic DNA from UV and iron-induced oxidative damage in certain cell types^43,45^. Ferritin may also scavenge other toxic metal species such as beryllium^46^, and together, these mechanisms may prevent germline mutations that may otherwise not be repaired until fertilization, if at all^47,48^.

The proteasome structures reported here revealed novel intermediates in PA200-mediated activation *in situ*. We identified that the PA200 HbYX motif binding to the α5/α6 pocket is a primary interaction, but, notably, is insufficient to open the α gate. Rather, additional interactions mediated by the PA200 α1/α2 loop, N-terminal extensions of α subunits, or other peripheral PA200 sites are necessary for full gate opening. While occupancy of the α1/α2 pocket is also a prominent feature of proteasome binding by the ATP-dependent 19S activator^49–52^, this interaction alone is insufficient to promote gate opening^53^, consistent with our finding that multiple interactions are required to open the 20S gate. Moreover, our structures reveal a highly dynamic association of PA200 with the 20S core, in which significant conformational flexibility is maintained until complete PA200 association and gate opening in the full-bound state (Figure 5). This mechanism may minimize non-specific proteolysis by shielding the 20S active sites until substrates are completely encapsulated within the PA200 cavity and proximal to the α gate.

In summary, our analysis provides novel insight into the molecular architecture of sperm nuclear vacuoles. These results provide a framework for studying the nuclear architecture of diverse cell types at high resolution in near-native environments.

## Acknowledgements

We thank Tyler Brittain for helpful discussions regarding cryo-ET data processing, Xiang Feng for assistance with tomogram denoising, and Pamela J. Bjorkman and David Chan for critical reading of the manuscript. J.R.B. is a Fellow of The Jane Coffin Childs Memorial Fund for Medical Research and of the Caltech Presidential Postdoctoral Fellows program. This project was supported by National Science Foundation Graduate Research Fellowship Program grants DGE-2146755 (to S.Q.) and DGE-1752814 and DGE-2146752 (to W.M.S.), a Pew Biomedical Scholars award (to P.V.L.), a grant from the Global Consortium for Reproductive Longevity and Equality (to P.V.L.), and a research grant from the Shurl and Kay Curci Foundation (to Z.C.). This paper was typeset with the bioRxiv Word template by @Chrelli: www.github.com/chrelli/bioRxiv-word-template.

## Author contributions

J.R.B. and Z.C. conceived the project and designed experiments. W.M.S. prepared the human sperm sample under the guidance of P.V.L. Z.C. prepared sperm EM grids. M.S. and N.J. generated sperm lamellae. J.R.B., R.Y., X.Z., and Z.C. collected TEM data. J.R.B. processed and analyzed TEM data. S.Q. performed unbiased AlphaFold docking. J.R.B. and S.Q. performed molecular modeling. J.R.B. and Z.C. wrote and edited the manuscript with inputs from all authors.

## Data availability

Cryo-EM densities have been deposited at the Electron Microscopy Data Bank (EMDB) under accession codes EMD-76923 (20S), EMD-76924 (20S-PA200 consensus), EMD-76925 (20S-PA200 full), EMD-76926 (20S-PA200 partial), EMD-76927 (20S-PA200 weak), EMD-76928 (20S-PA200 double), and EMD-76929 (ferritin). Atomic coordinates have been deposited at the Protein Data Bank (PDB) under accession codes PDB 13AT (20S), PDB 13AU (20S-PA200 consensus), PDB 13AV (20S-PA200 full), PDB 13AW (20S-PA200 partial), and PDB 13AX (20S-PA200 weak).

## Methods

### Sample collection

A freshly ejaculated semen sample was obtained by masturbation from a man aged 25-39 years old. The sample was verified by light microscopy to be normozoospermic with a cell count of at least 30 million sperm per milliliter. Sperm were isolated using the swim-up procedure, as described previously^54^. All experimental procedures using human-derived samples were approved by the Committee on Human Research at the University of California, Berkeley, under IRB protocol number 2013-06-5395.

Mouse sperm were collected from 10 to 16-week-old C57BL/6J mice following a published protocol^55^. Briefly, the sperm were extracted from vasa deferentia by applying pressure to cauda epididymides in Krebs buffer (1.2 mM KH_2_PO_4_, 120 mM NaCl, 1.2 mM MgSO_4_·7H_2_O, 14 mM dextrose, 1.2 mM CaCl_2_, 5 mM KCl, 25 mM NaHCO_3_). The sperm were washed and resuspended in ∼100 µL Krebs buffer for the following experiments.

### Grid preparation

EM grids (Quantifoil R 2/2 Au 200 mesh) were glow-discharged using an easiGlow system (PELCO). Grids were then loaded onto a Leica GP2 plunge freezer (pre-equilibrated to 95% relative humidity at 25 °C). 3.5 µL of the sperm mixtures were loaded onto each grid, followed by a 15-second incubation period. The grids were then blotted for 4 seconds and plunge-frozen in liquid ethane cooled by liquid nitrogen.

### Cryogenic focused ion beam (cryo-FIB) milling

Cryo-FIB milling was performed using an Aquilos 2 cryo-FIB/SEM microscope (Thermo Fisher Scientific). Panorama SEM maps of entire grids were first taken at 377x magnification using an acceleration voltage of 2 kV with a beam current of 13 pA, and a dwell time of 1 µs. Targets with appropriate thickness for milling were selected on each grid. For all grids, an inorganic platinum layer (∼10 nm) was deposited by sputter coating; a gas injection system (GIS) was then used to deposit the compound trimethyl(methylcyclopentadienyl)platinum(IV), followed by a final inorganic platinum layer. The stage was tilted to 8°, corresponding to a milling angle of 11° relative to the plane of the grids. FIB milling was performed using stepwise decreasing current as the lamellae became thinner (0.5 nA to 30 pA, final thickness: ∼150 nm). The grids were then stored in liquid nitrogen before data collection.

### Image acquisition and tomogram reconstruction

Tilt series of human and mouse sperm nuclei lamellae were collected on a Titan Krios transmission electron microscope (Thermo Fisher Scientific) operated at 300 kV and equipped with a high brightness field emission gun (xFEG), a spherical aberration corrector, and a BioQuantum K3 Imaging Filter (Gatan) with a zero-loss energy slit width between 6 and 10 eV. Images were recorded as 20-frame movies at a nominal magnification of 42,000x in super-resolution counting mode with correlated double sampling using SerialEM^56^, corresponding to a nominal physical pixel size of 1.71 Å. For each tilt series, images were acquired using a modified dose-symmetric scheme between -40° and 40° relative to the lamella, with 4° increments and groupings of two images on each side (0°, 4°, 8°, -4°, -8°…), using a defocus range of -2 to -4 µm. The total electron dose per tilt series was 120 e^-^/Å^2^, applied at a dose rate of ∼13 e^-^ per pixel per second.

All movies were gain-corrected and corrected for global drift, Fourier-cropped by a factor of 2 (to achieve the physical pixel size, 1.71 Å), and summed without dose weighting using MotionCor2^57^. Tilt series alignment and tomogram reconstruction of mouse data were performed in Are-Tomo2^58^ with no further processing. For human data, frame CTF estimation was performed in WarpTools^59^, followed by tilt series alignment in AreTomo2, and tilt CTF estimation and tomogram reconstruction (bin 8, 13.68 Å/pix) in WarpTools. Tomograms were visually inspected, and 111 (from an initial 776 complete tilt series collected) were selected for further processing.

### Particle picking and sub-tomogram averaging

In all automated picking schemes, we sought to substantially over-pick to maximize particle number, followed by duplicate removal to generate the final stacks. The initial particle numbers listed in Table S1, representing the number used as inputs to initial 3D classification jobs, are thus inflated by false-positives and duplicates. This strategy resulted in the recovery of the majority of particles for all species (Figure 1A,B).

Proteasome particles were picked using TomoTwin^60^ using five references corresponding to different particle orientations. Duplicate particles were removed within 100 Å, and were then cleaned by two rounds of 2D classification in RELION-4^31^ (bin 8, 13.68 Å/pix, 22 pix box, K=100, T=1). Particles were extracted as 2D image series in WarpTools (bin 4, 6.84 Å/pix, 64 pix box), and further cleaned by three rounds of 3D classification in RELION-5^61^ (K=3-5, T=4), yielding two maps, corresponding to a 20S proteasome and a 20S proteasome with additional density at one end. Initial 3D refinement was performed in RELION-5 using particles extracted at bin 2 (3.42 Å/pix, 128 pix box), applying *C*2 symmetry to the 20S particle and *C*1 symmetry to the map with additional density. The resulting alignments were imported into M^16^, and multiple rounds of refinement were performed, progressively adding refinement of 2D image warp, particle poses, defocus, stage angles, anisotropic magnification, spherical aberration, and third-order Zernike polynomials. A final round of duplicate removal was performed using the refined particle centers (within 100 Å).

The cleaned proteasome stack with additional density was then subjected to three classification schemes to focus on various aspects of the observed heterogeneity, followed by a final series of M refinements with the stacks corresponding to the 20S proteasome and high-resolution ferritin map (see below). Particles were re-extracted in WarpTools (bin 2, 3.42 Å/pix, 152 pix box), and subjected to further 3D classification in RELION-5 without image alignment. A mask above the *trans* 20S α ring (without additional density visible) was used for focused classification to generate the single- and double-bound maps (K=5, T=4). A mask around the *cis* 20S α ring and additional density was used for focused classification to isolate the highest-quality stack (K=4, T=4), which was refined to generate the consensus high-resolution map with additional density. The same mask was used to obtain the three bound intermediates (supervised classification with maps from a prior round of classification, K=3, T=4).

Non-proteasomal spherical particles were picked manually and by crY-OLO^62^, using two models, one trained on all such particles and one excluding particles with strong central density. Particles were extracted in Warp-Tools (bin 4, 6.84 Å/pix, 64 pix box) and subjected to 3D classification in RELION-5 (K=10, T=4). Particles from a class resembling a hollow sphere were re-extracted at 4 Å/pix (64 pix box) and refined with *O* symmetry after duplicate removal within 130 Å. These particles were imported into the M project used to refine the proteasome maps (after final classifications) and subjected to the same rounds of multi-particle refinement as described above. To obtain a map that contained the strong central density observed in tomograms, manually picked particles (with and without visible central density) were extracted in WarpTools (4 Å/pix, 64 pix box), subjected to 3D classification (K=2, T=4, *O* symmetry) to separate the two populations, and refined in RELION-5 with *O* symmetry.

Following processing, the pixel size of the data was refined to 1.68 Å by calculating correlation values between the maps and previously determined structures in UCSF ChimeraX^63^. Map B factors and gold-standard resolutions (Fourier shell correlation between two independently-refined half-maps, threshold of 0.143) were calculated in RELION-5 using the calibrated pixel size. Local resolution maps were calculated in M.

### Unbiased structural identification by global density matching

To identify candidate protein structures corresponding to the observed densities, an unbiased global docking procedure was performed using the COLORES program from the Situs package^5,64,65^. A total of 23,577 human protein structures from the AlphaFold Protein Structure Database^14,15^ were docked against the experimental density maps.

For the proteasome-associated density, the map resolution used for docking was 4.6 Å. The density was segmented using UCSF ChimeraX, and a density threshold cutoff of 0.003 was selected based on visual inspection of the density distribution. Rigid-body docking was performed with an angular search step of 10°. After exhaustive matching against the AlphaFold database, candidate structures were ranked based on the correlation coefficient. PA200 yielded the highest correlation score among all screened structures, and the resulting docked model showed clear structural agreement with the density, including secondary structures throughout the majority of the sequence, supporting the identification of the density as PA200.

For the ferritin density, a single monomeric subunit was segmented from the map in UCSF ChimeraX to generate the search target. The docking resolution was set to 8.0 Å with a density cutoff of 0.002. Because the small box size of the ferritin subunit could lead to artificially inflated correlation coefficients (>1) during correlation evaluation, the search volume was expanded to approximately three times the size of the segmented density by padding the map boundaries and modifying corner voxels to prevent cropping in COLORES. Rigid-body docking was performed with an angular search step of 10°. The resulting correlation scores were ranked across all screened AlphaFold models. Examination of the top 20 scoring models revealed that ferritin subunits consistently ranked among the highest matches, together with structurally related proteins sharing the characteristic four-helix bundle topology, confirming the assignment of the density as ferritin.

### *PSMA7*/*PSMA8* assignment with Rosetta

To determine the identity of the α4 subunit in vacuolar proteasomes, Rosetta density-guided relaxation against the 20S map (locally-filtered) was performed using the FastRelax protocol^66,67^. Initial atomic models of *PSMA7* and *PSMA8* gene products (AlphaFold3 predictions, canonical *PSMA7* sequence and *PSMA8* splice variant Q8TAA3-5) were manually docked into the segmented subunit density using UCSF ChimeraX, ensuring identical backbone placement for both models prior to refinement. Rosetta density relax simulations were then performed under varying density-weighting parameters to evaluate how strongly the density map influences the structural refinement. The weight applied to the density score term (*elec_dens_fast*) was varied from 20 to 35 in increments of 5. For each density weight, 10 independent refinement runs were performed to account for stochastic sampling during the Rosetta relaxation procedure. The resulting models were evaluated using EM_Match score, which measures the correlation between the refined structure and the density map, and Rosetta total energy (REU). Inspection of results shows consistently superior fits for *PSMA8*, suggesting that the density corresponds to this paralog (Figure S6D). *PSMA8* was therefore used to build all proteasome models.

### Molecular modeling

To model the sidechain density of the 20S structure, *PSMA7* was substituted with the AlphaFold2 prediction of *PSMA8* (Q8TAA3-5) in a previously published model (PDB 7NAN^20^), residues outside the density were removed in Coot^68^, and Rosetta density-guided relaxation was performed using the FastRelax protocol (torsion-space refinement, *elec_dens_fast*=35), with C2 symmetry imposed^66,67,69,70^. To model the 20S-PA200 consensus structure, *PSMA8* was replaced as above in a previously published model (PDB 6KWY^19^), Coot was used to remove residues outside the density, and Rosetta FastRelax was used (torsion-space refinement, *elec_dens_fast*=20) to relax into the locally-filtered density. These models, or portions thereof, were used as starting points to model the full, partial, and weak 20S-PA200 structures in Rosetta and ISOLDE^71^. The 20S-PA200 weak model was generated solely by rigid-body docking chains from the 20S model into the corresponding map in Phenix^72^. Residues in all models except that of the 20S proteasome were truncated to alanine due to the moderate map resolutions. Estimation of model *B* factors was performed in Phenix, and Coot and ISOLDE were used to finalize all models. All references to proteasome α and β subunits follow yeast nomenclature (for example, gene product of *PSMA7* = α4).

### Calculation of particle distance to vacuole edge

The distances of proteasome and ferritin particles to the vacuole edge were measured in 10 tomograms in which the vacuole-chromatin boundary was visible in all directions in the xy plane, representing ∼40% of all identified ferritin particles. Geometric models of vacuoles were generated in ArtiaX^73^ using a boundary around all particles in each vacuole (final poses after M refinement). Euclidean distances of each particle to the nearest voxel of the vacuole sidewall were calculated using a custom Python script. Significance testing for the ferritin and proteasome distance distributions was performed using Welch’s two-tailed t-test.

### Data analysis and figure preparation

Electron microscopy maps and models were rendered in UCSF ChimeraX, and tomogram slices were rendered in IMOD^74^. 3D visualization of particles in tomograms was performed in UCSF ChimeraX using the ArtiaX plugin. Where applicable, tomogram denoising was performed using CryoSamba^75^. Numerical data were analyzed and plotted using GraphPad Prism. Figures were prepared using Adobe Illustrator. Software used in this project was installed and configured by SBGrid^76^.

---

**Figure S1.**
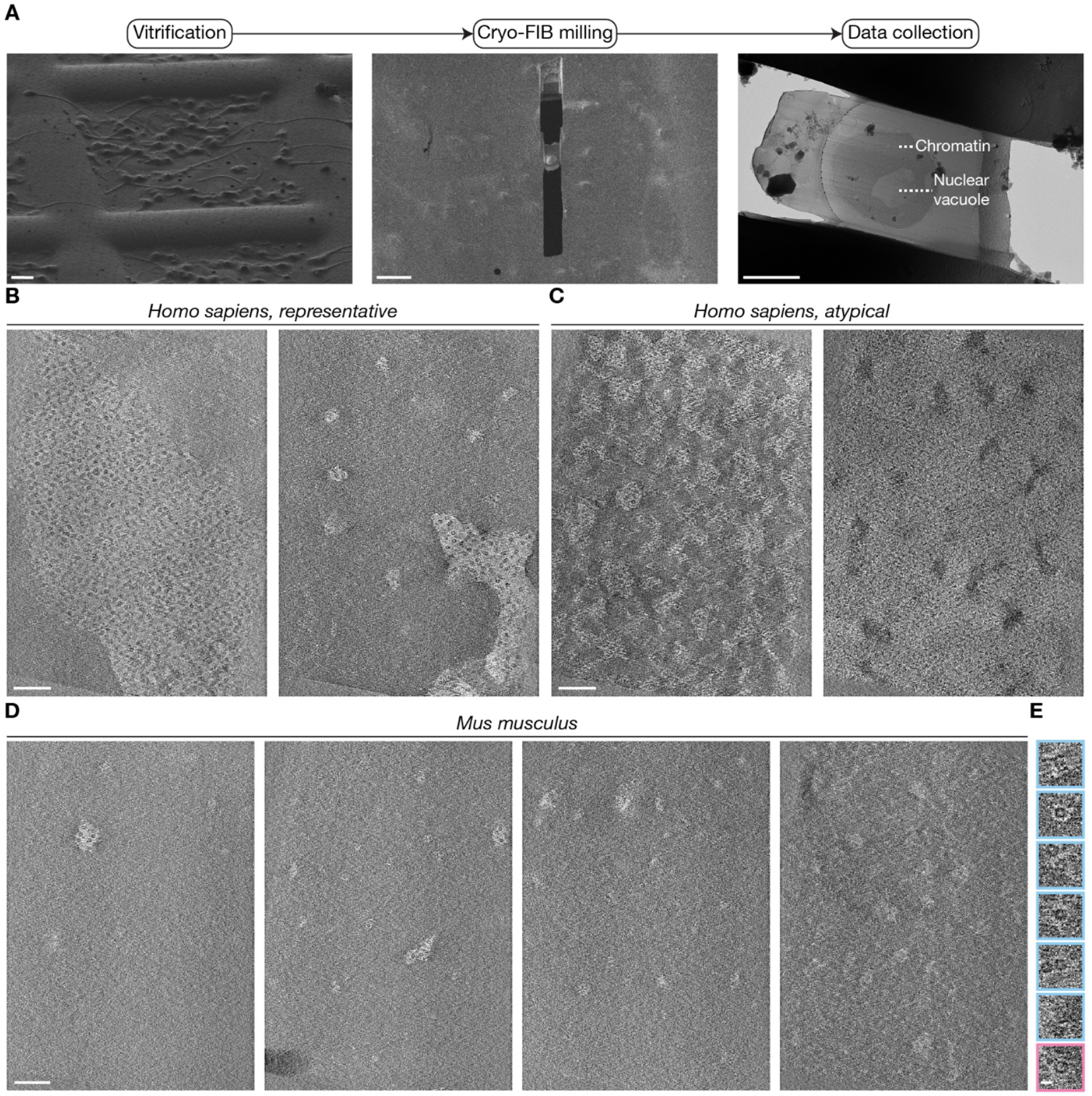
Cryo-ET of mammalian sperm nuclei reveals conserved vacuole morphology and protein complexes. **(A)** Representative images illustrating the cryo-focused ion beam milling and transmission electron microscopy workflow for mammalian sperm nuclei. Images of human sperm are shown. **(B)** Representative z-slices of tomograms of human sperm nuclei with normal vacuole morphology. **(C)** Representative z-slices of tomograms of human sperm nuclei with non-canonical morphology, including interconnected networks of vacuole-like features and seemingly decondensed chromatin. **(D)** Representative z-slices of tomograms of mouse sperm nuclei. **(E)** Gallery of individual particles identified in mouse sperm nucleus tomograms (central z-slices). Particles outlined in blue correspond to proteasomes; the particle outlined in pink corresponds to a ferritin molecule. Scale bars, 10 μm, 10 μm, and 2 μm **(A)**, 100 nm **(B, C, D)**, and 10 nm **(E)**.

**Figure S2.**
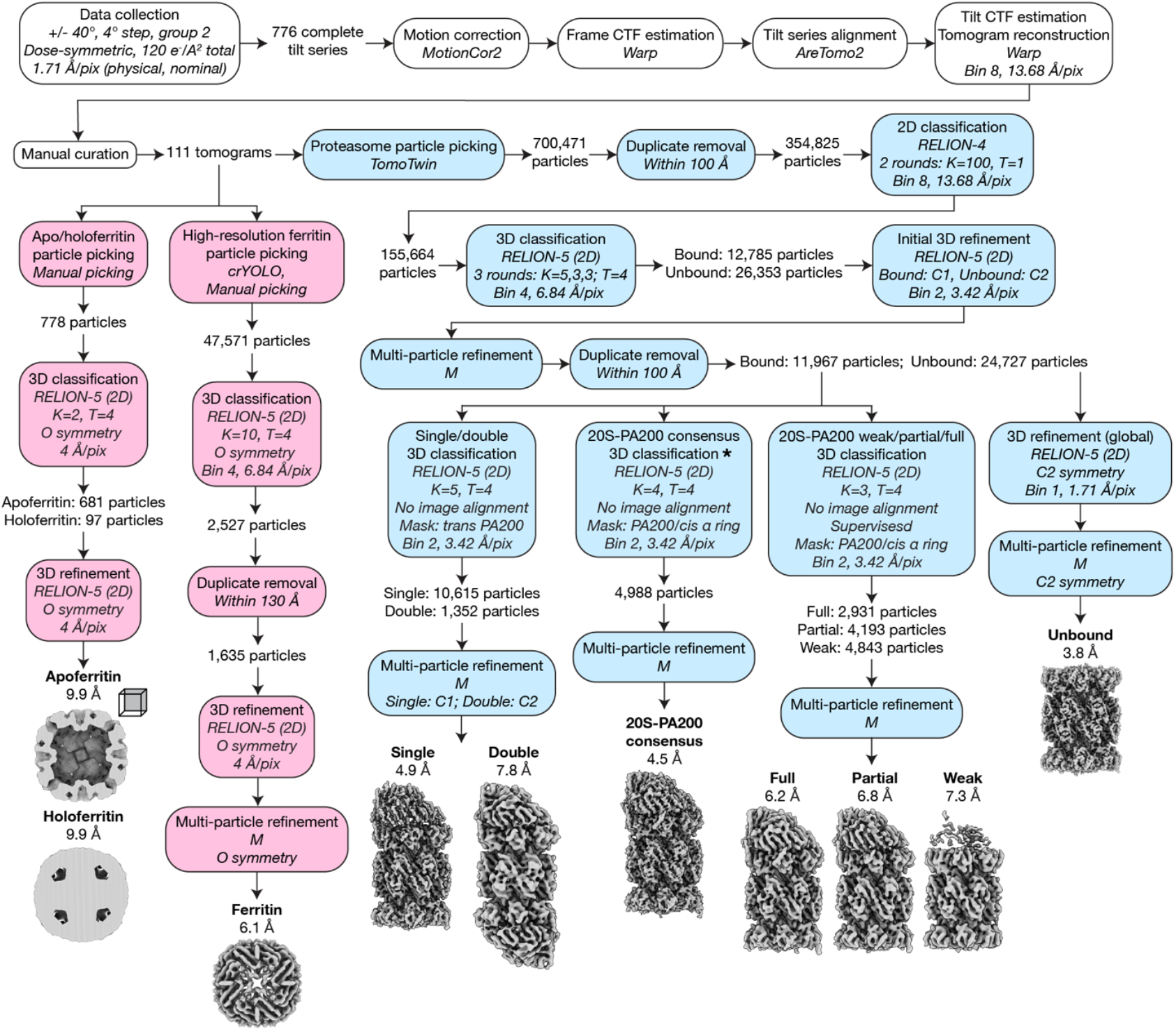
Human sperm nucleus high-resolution cryo-ET processing pipeline. Workflow for data collection and sub-tomogram averaging of proteasome and ferritin structures from human sperm nuclear vacuoles. Proteasome processing steps are colored blue, ferritin steps in pink. The 3D classification job described in Figure S6 is indicated (*).

**Figure S3.**
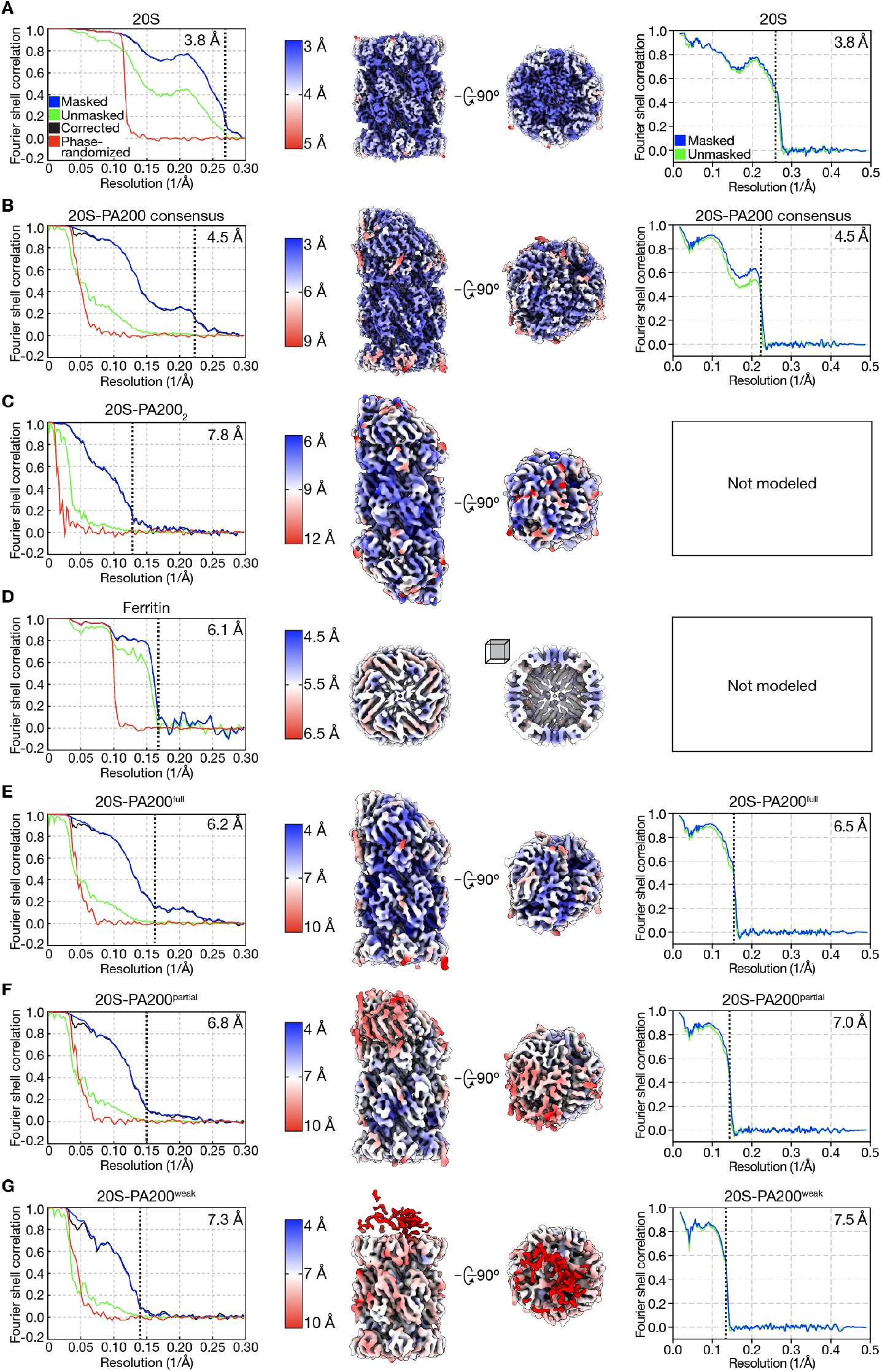
Cryo-ET map and model validation. Gold-standard Fourier shell correlation (FSC) curves, filtered maps colored by local resolution, and map-model FSC curves for deposited models (where applicable) for **(A)** 20S, **(B)** 20S-PA200 consensus, **(C)** 20S-PA200_2_, **(D)** ferritin, **(E)** 20S-PA200^full^, **(F)** 20S-PA200^partial^, and **(G)** 20S-PA200^weak^. Displayed map resolutions are calculated from corrected curves (cutoff = 0.143), and displayed model resolutions are calculated from masked curves (cutoff = 0.5).

**Figure S4.**
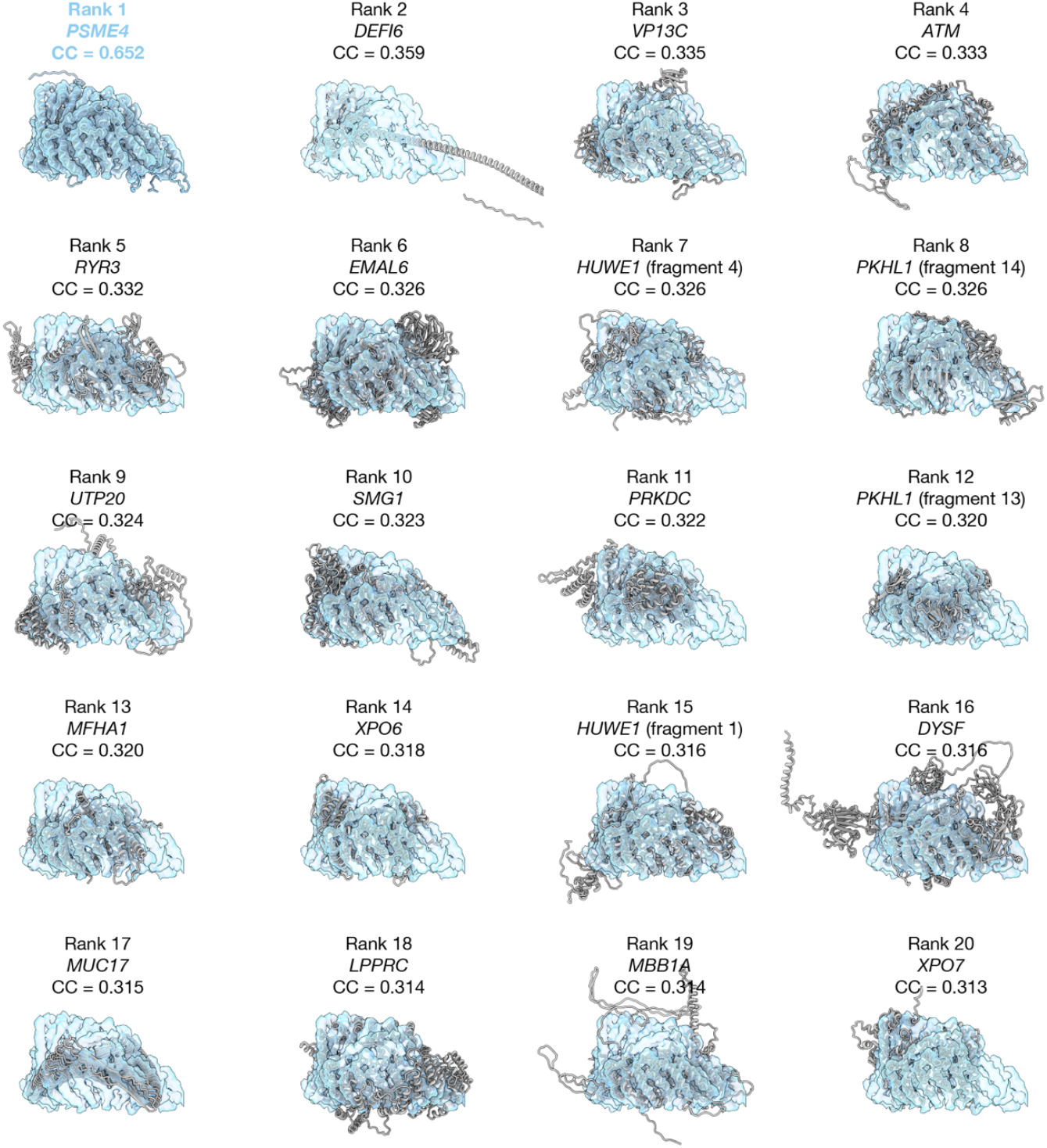
AlphaFold docking into the proteasome-associated density. The top 20 models from docking of the human AlphaFold proteome into the proteasome-associated density, overlaid with the map used. Rank, gene names, and cross-correlation scores are shown. Model of the best fit, PA200 (gene name *PSME4*) is colored blue, others in gray.

**Figure S5.**
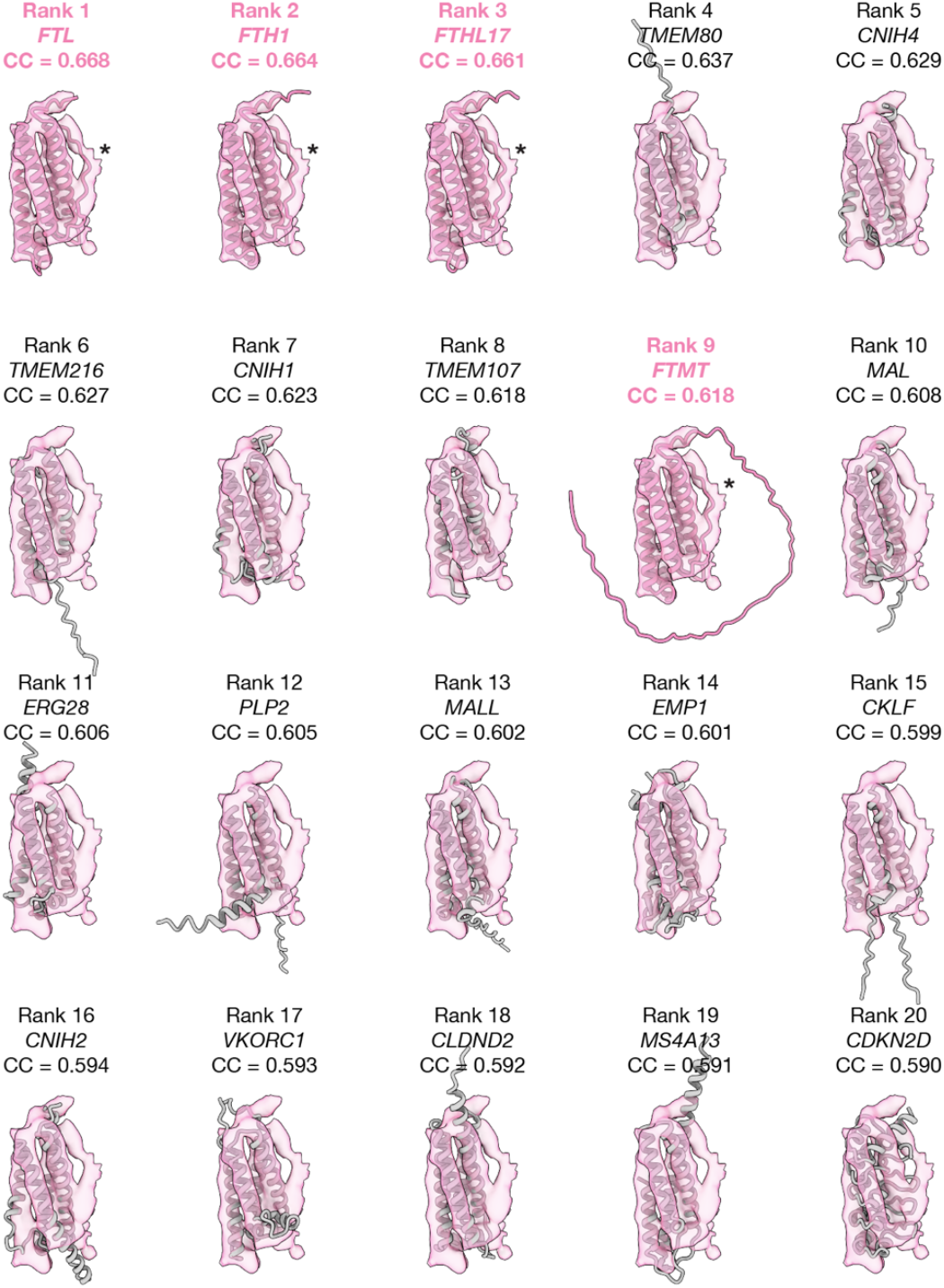
AlphaFold docking into the ferritin asymmetric unit. The top 20 models from docking of the human AlphaFold proteome into the ferritin asymmetric unit, overlaid with the map used. Rank, gene names, and cross-correlation scores are shown. Models of ferritin homologs are colored pink, others in gray. An ordered loop that only ferritin homologs contain is indicated in the ferritin fits (*).

**Figure S6.**
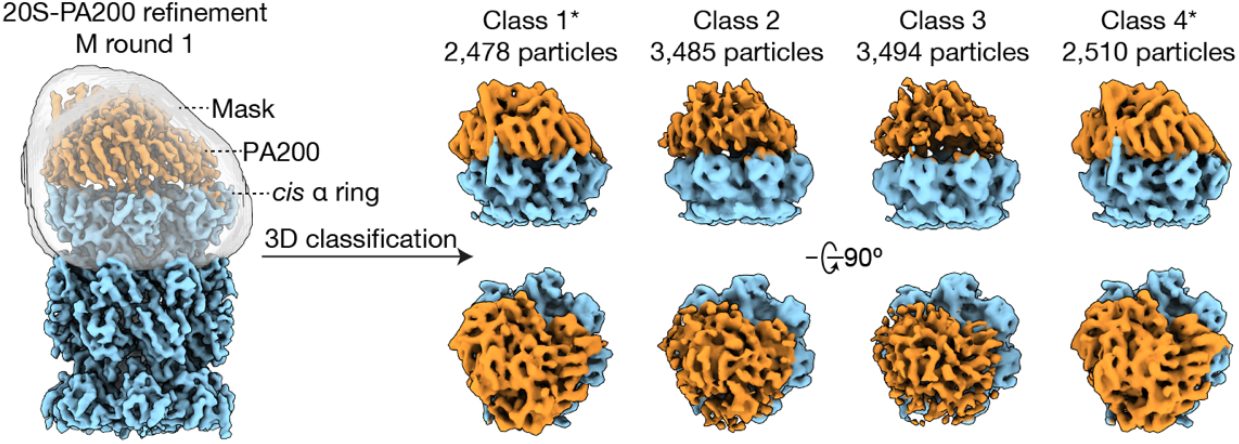
Conformational variability in 20S-PA200 complexes *in situ*. Schematic of 3D focused classification job used to generate the 20S-PA200 consensus map, colored as in Figure 2 (see Figure S2, *). The mask used is colored in transparent gray, overlaid with the reconstruction of the input particles. The output maps are displayed at thresholds such that the 20S density appears consistent between classes. Classes used in the final consensus refinement are indicated (*).

**Figure S7.**
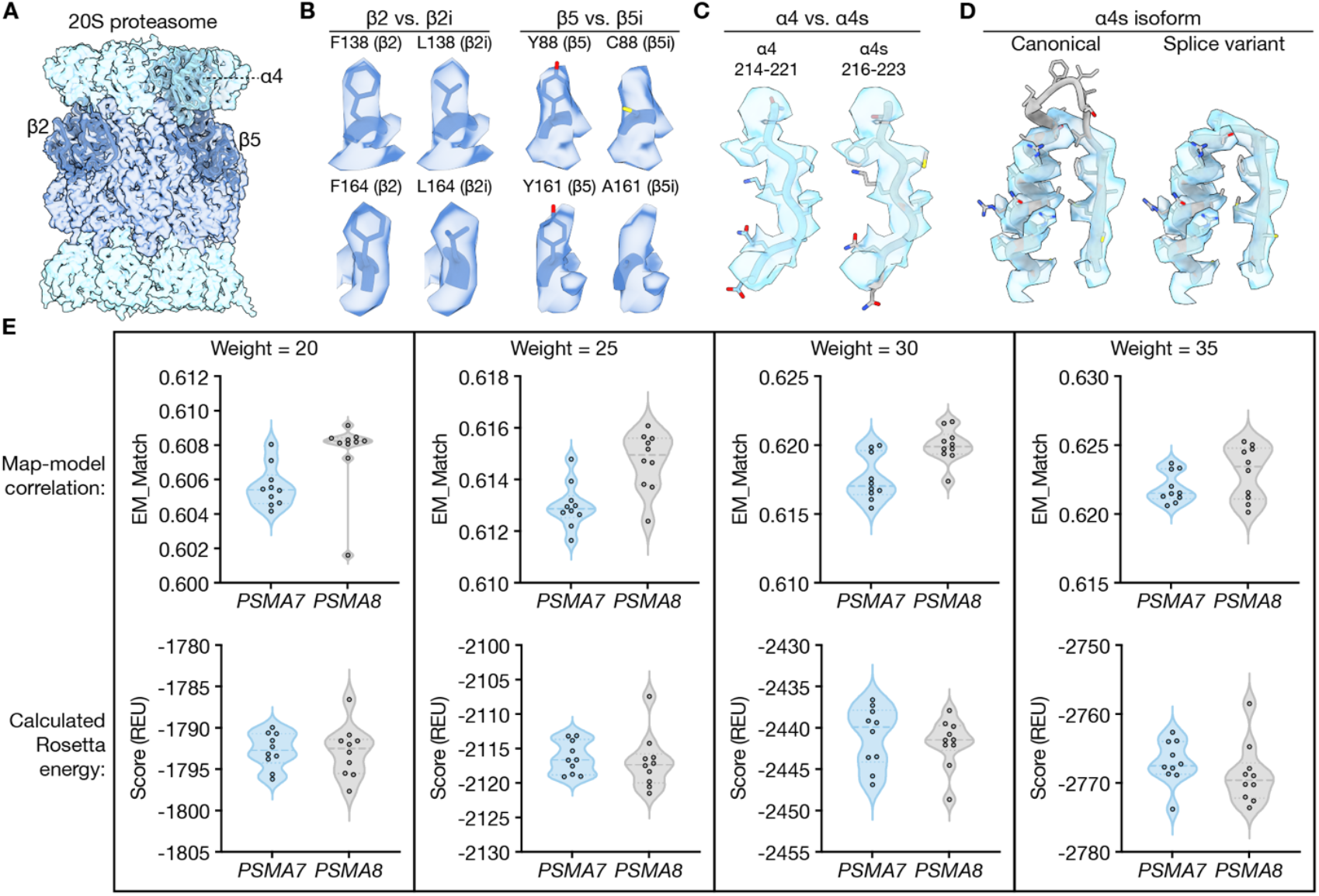
*In situ* proteasome subunit paralog determination. **(A)** 20S map (transparent) with models of α4, β2, and β5 shown. **(B)** Views of amino acid substitutions between constitutive and immunoproteasome subunits (constitutive, 20S model; immunoproteasome, PDB 6E5B). **(C)** Overlays of α4 (20S model, residues 214-221) and α4s (AlphaFold3 prediction, residues 216-223, Q8TAA3-5 numbering) on the 20S map (locally filtered), showing multiple amino acid substitutions. The identity at any position cannot be determined definitively. **(D)** Comparison of AlphaFold3 predictions of two splice isoforms of *PSMA8* (encoding α4s) docked into the 20S map (locally filtered). The canonical isoform (accession Q8TAA3-1, residues 71-99 shown) has a six-residue insertion incompatible with the density, while an alternatively spliced isoform (accession Q8TAA3-5, residues 71-93 shown) does not. **(E)** Violin plots of Rosetta density relax EM_Match and REU scores for *PSMA7* and *PSMA8* (splice variant) models, for various map weights (Methods). Medians (thick dashes) and quartiles (thin dashes) are shown, overlaid with individual values. Higher EM_Match and lower REU scores for *PSMA8* indicate a better fit for this paralog.

**Figure S8.**
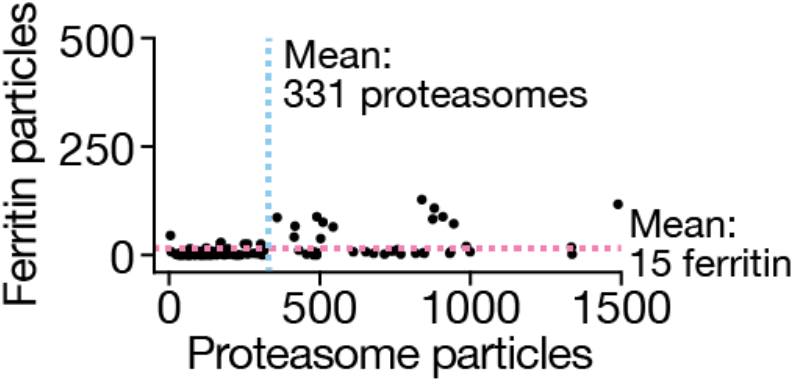
Ferritin and proteasome counts in human tomograms. Ferritin vs. proteasome counts in human sperm nuclear vacuole tomograms (maximum field of view ∼0.7 μm^2^), including all particles used in final reconstructions (*n* = 111 tomograms). Mean values for each distribution are shown.

**Figure S9.**
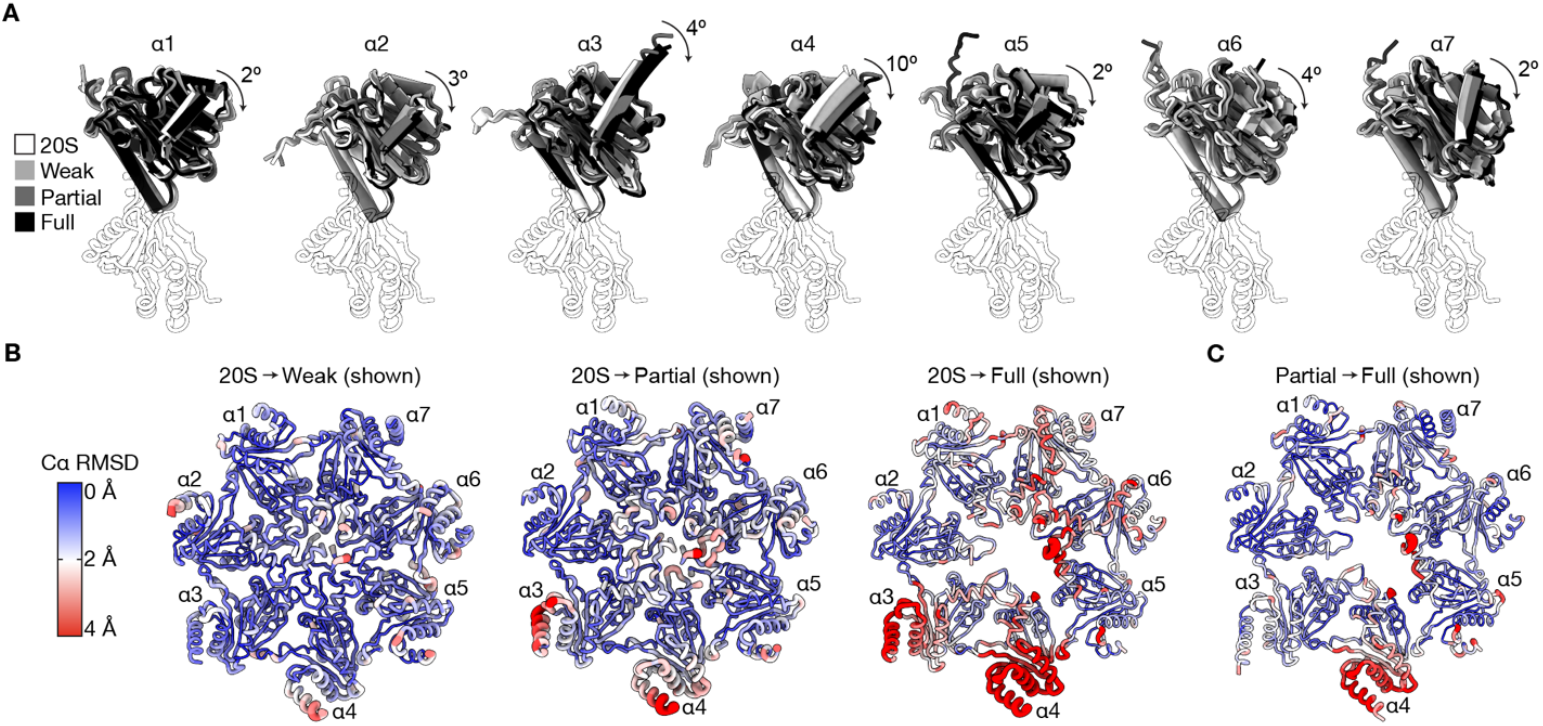
Conformational changes in 20S-PA200 complexes *in situ*. **(A)** Overlay of α subunits of 20S and *cis* 20S-PA200 complexes, aligned by the underlying β subunit. The α4 overlay is as in Figure 4D. Angles of rotation between 20S and 20S-PA200^full^ are shown for each α subunit. **(B)** *Cis* α gates of the weak, partial, and full 20S-PA200 structures, colored by Cα r.m.s.d. values relative to the 20S structure. **(C)** *Cis* α gates of the 20S-PA200 full structure, colored by Cα r.m.s.d. values relative to the partial structure, as in **(B)**.

**Table S1.** Cryo-ET data collection, refinement, and validation statistics.

|  | 20S (EMD-76923, PDB 13AT) | 20S-PA200 consensus (EMD-76924, PDB 13AU) | 20S-PA200<sup>full</sup> (EMD-76925, PDB 13AV) | 20S-PA200<sup>partial</sup> (EMD-76926, PDB 13AW) | 20S-PA200<sup>weak</sup> (EMD-76927, PDB 13AX) | 20S-PA200<sub>2</sub> (EMD-76928) | Ferritin (EMD-76929) |
| --- | --- | --- | --- | --- | --- | --- | --- |
| **Data collection and processing** |  |  |  |  |  |  |  |
| Microscope and camera | Titan Krios, K3 | Titan Krios, K3 | Titan Krios, K3 | Titan Krios, K3 | Titan Krios, K3 | Titan Krios, K3 | Titan Krios, K3 |
| Magnification | 42,000 | 42,000 | 42,000 | 42,000 | 42,000 | 42,000 | 42,000 |
| Voltage (kV) | 300 | 300 | 300 | 300 | 300 | 300 | 300 |
| Data acquisition software | SerialEM | SerialEM | SerialEM | SerialEM | SerialEM | SerialEM | SerialEM |
| Electron exposure (e-/Å<sup>2</sup>) | 120 | 120 | 120 | 120 | 120 | 120 | 120 |
| Tilt range/increments (°) | ±40/4 | ±40/4 | ±40/4 | ±40/4 | ±40/4 | ±40/4 | ±40/4 |
| Defocus range (µm) | -2 to -4 | -2 to -4 | -2 to -4 | -2 to -4 | -2 to -4 | -2 to -4 | -2 to -4 |
| Pixel size (Å, calibrated) | 1.68 | 1.68 | 1.68 | 1.68 | 1.68 | 1.68 | 1.68 |
| Symmetry imposed | C2 | C1 | C1 | C1 | C1 | C2 | O |
| Initial particle images (no., see Methods) | 155,664 | 155,664 | 155,664 | 155,664 | 155,664 | 155,664 | 47,571 |
| Final particle images (no.) | 24,727 | 4,988 | 2,931 | 4,193 | 4,843 | 1,352 | 1,635 |
| Map resolution (Å) | 3.8 | 4.5 | 6.2 | 6.8 | 7.3 | 7.8 | 6.1 |
| FSC threshold | 0.143 | 0.143 | 0.143 | 0.143 | 0.143 | 0.143 | 0.143 |
| Map resolution range (Å) | 3.5-5 | 3.5-10 | 4-12 | 5-15 | 6-20 | 5-12 | 5-6 |
| **Refinement** |  |  |  |  |  |  |  |
| Initial model used (PDB, AlphaFoldDB accession) | 7NAN, AF-Q8TAA3-5-F1 | 6KWY, AF-Q8TAA3-5-F1 | 6KWY, AF-Q8TAA3-5-F1 | 7NAN, 6KWY, AF-Q8TAA3-5-F1 | 7NAN, AF-Q8TAA3-5-F1 | N/A | N/A |
| Model resolution (Å) | 3.8 | 4.5 | 6.5 | 7.0 | 7.5 | N/A | N/A |
| FSC threshold | 0.5 | 0.5 | 0.5 | 0.5 | 0.5 | N/A | N/A |
| Map sharpening *B* factor (Å<sup>2</sup>) | -76.8 | -17.1 | -32.3 | -37.5 | -186.0 | -139.4 | -344.8 |
| **Model composition** |  |  |  |  |  |  |  |
| Nonhydrogen atoms | 48,222 | 38,943 | 39,052 | 39,174 | 30,476 | N/A | N/A |
| Protein residues | 6,192 | 7,900 | 7,922 | 7,947 | 6,192 | N/A | N/A |
| ***B* factors (Å<sup>2</sup>)** |  |  |  |  |  |  |  |
| Protein | 65.96 | 41.54 | 115.27 | 192.97 | 259.42 | N/A | N/A |
| **R. m. s. deviations** |  |  |  |  |  |  |  |
| Bond lengths (Å) | 0.01 | 0.003 | 0.003 | 0.003 | 0.006 | N/A | N/A |
| Bond angles (°) | 0.783 | 0.666 | 0.668 | 0.671 | 0.821 | N/A | N/A |
| **Validation** |  |  |  |  |  |  |  |
| MolProbity score | 0.98 | 1.15 | 1.63 | 1.63 | 0.66 | N/A | N/A |
| Clashscore | 2.09 | 0.75 | 3.92 | 4.77 | 0.47 | N/A | N/A |
| Poor rotamers (%) | 0.04 | N/A | N/A | N/A | N/A | N/A | N/A |
| **Ramachandran plot** |  |  |  |  |  |  |  |
| Favored (%) | 98.56 | 93.92 | 94.40 | 94.40 | 98.58 | N/A | N/A |
| Allowed (%) | 1.34 | 6.02 | 5.47 | 5.47 | 1.32 | N/A | N/A |
| Disallowed (%) | 0.10 | 0.06 | 0.13 | 0.13 | 0.10 | N/A | N/A |

